# Genome-scale host-pathogen prediction for non-medical microbes

**DOI:** 10.1101/2022.07.20.500869

**Authors:** Mais Ammari, Cathy Gresham, Fiona M McCarthy, Bindu Nanduri

## Abstract

**Background:** Network studies of host-pathogen interactions (HPI) are critical in understanding the mechanisms of pathogenesis. However, accessible HPI data for agriculturally important pathogens are limited. This lack of HPI data impedes network analysis to study agricultural pathogens, for preventing and reducing the severity of diseases of relevance to agriculture.

**Results:** To rapidly provide HPIs for a broad range of pathogens, we use an interolog-based approach. This approach uses sequence similarity to transfer known HPIs from better studied host-pathogen pairs and predicts 389,878 HPIs for 23 host-pathogen systems of relevance to US agriculture. Each predicted HPI is qualitatively assessed using co-localization, infection related processes, and interacting domains and this information is provided as a confidence indicator for the prediction. Evaluation of predicted HPIs demonstrates that the host proteins predicted to be involved in pathogen interactions include hubs and bottlenecks in the network, as reported in curated host proteins. Moreover, we demonstrate that the use of the predicted HPIs adds value to network analysis and recapitulates known aspects of host-pathogen biology. Access to the predicted HPIs for these agricultural host-pathogen systems is available via the Host Pathogen Interaction Database (HPIDB, hpidb.igbb.msstate.edu), and can be downloaded in standard MITAB file format for subsequent network analysis.

**Conclusions:** This core set of interolog-based HPIs will enable animal health researchers to incorporate network analysis into their research and help identify host-pathogen interactions that may be tested and experimentally validated. Moreover, the development of a larger set of experimentally validated HPI will inform future predictions. Our approach of transferring biologically relevant HPIs based on interologs is broadly applicable to many host-microbe systems and can be extended to support network modeling of other pathogens, as well as interactions between non-pathogenic microbes.

## BACKGROUND

Host-pathogen interactions (HPI) dictate the course of infection and pathogenesis and understanding these interactions provides the fundamental basis for rational design of novel therapeutic strategies. The rapid growth of HPI data has led to the application of network analysis approaches in human health and disease research (1,2). However, few HPIs related to pathogens relevant to agriculture species are available in molecular interaction databases. This lack of accessible HPI data hinders the application of network analyses to better understand and prevent and/or reduce the severity and economic impact of diseases of livestock and aquaculture. Likewise, the HPI data available from public interaction databases is limited to a small set of hosts and their pathogens. Therefore, there is a need for a broad range of HPI data for agricultural host-pathogen systems.

Given the large number of infectious diseases relevant to livestock species, genome-scale identification of HPI using high throughput experimental approaches can be both expensive and time consuming. Even for a single host such as bovine, validating HPI with multiple pathogens could be impractical given the large interaction space i.e., ~22,000 bovine genes (3,4) and their interaction with thousands of pathogen proteins. Computational prediction of HPI is a pragmatic approach to rapidly generate HPI data for agricultural host-pathogen systems. A review of computational HPI prediction methods is available in literature (5,6). There are many studies describing HPI prediction for a single host pathogen pair or a single host and a pathogen group of pathogens i.e., bacteria, virus, or fungi (7–9). Computational prediction methods can build on experimental HPI, where available, or utilize protein information from experimental HPI to infer or predict interactions. The interolog method is one such approach to infer interactions between two proteins for any given host-pathogen system, based on experimental HPI from another host-pathogen pair (1,10). The basis of the interolog method is the conservation of the interaction experimentally verified in one system in the test host-pathogen system. HPI predictions by domain-domain interactions (11,12), domain-motif interactions (13), and structure (14), are augmented by network topology measures (15) of interacting proteins. Machine learning based approaches including multitask learning (16), multi-layer neural networks (29945178), when combined with network inference can be powerful tools for HPI prediction (17). Irrespective of the method used for prediction, qualitative assessment of predicted HPI routinely uses Gene Ontology (GO) annotations of the interacting pairs.

To enable access to HPIs that underpin health and disease in agriculturally important species, we provide curated HPI data via the HPIDB database (18). Although manual curation from published experimental data remains the ‘gold standard’ for HPI, there is limited or no experimental evidence available describing the HPI of many economically important agricultural pathogens. To supplement the existing HPI information and support functional modeling and hypothesis testing for animal studies, we utilize computational workflows for providing predicted HPI. We generate predicted HPI using interologs (19), which are conserved interactions between two proteins that have homologs in other species. Evolutionary conservation allows the transfer of the interaction from the original species to other species. Expansion of this concept to include HPI has been applied to human HPI data (20,21).

Using interolog method, we report the prediction of 389,878 HPIs for 23 host-pathogen pairs. Since we inferred these interactions based on experimental HPI, there is no definitive true positive or false negative information to assess the transfer of interaction. Therefore, we conducted aa qualitative assessment of interolog based inference by evaluating the biological features such as protein domains, motifs and structure of predicted HPI, combined with GO annotations, and assigned confidence indicators. We also demonstrate the use of predicted HPI in functional modeling of animal health data, by including network topology measures such as hubs and bottleneck proteins represented in the predicted HPI dataset. Our predicted interologs are freely and publicly available via HPIDB (hpidb.igbb.msstate.edu). This approach demonstrates the utility of inferring biologically relevant HPI that can be applied to many other types of important pathogens.

## METHODS

### Selecting host-pathogen pairs for predicting HPI using interolog approach

We prioritized the host-pathogen systems for HPI predictions by creating an initial list of pathogens that constitute priority pathogens identified by USDA stakeholders (22), active animal health grants funded through USDA, and existing HPI data and available HPI literature. Pathogens on our priority list cause disease in USDA livestock and aquaculture species and we have attempted to ensure that all important hosts are represented, as are classes of pathogens (gram-negative and gram-positive bacteria, RNA and DNA viruses, and parasites). Another consideration for selection of hosts and pathogens was ensuring a variety of systems to maximize the transfer of manual curation data to other host-pathogen systems using computational methods. A summary of the host-pathogen pairs chosen for analysis is shown in Table 1.

**Table 1.**
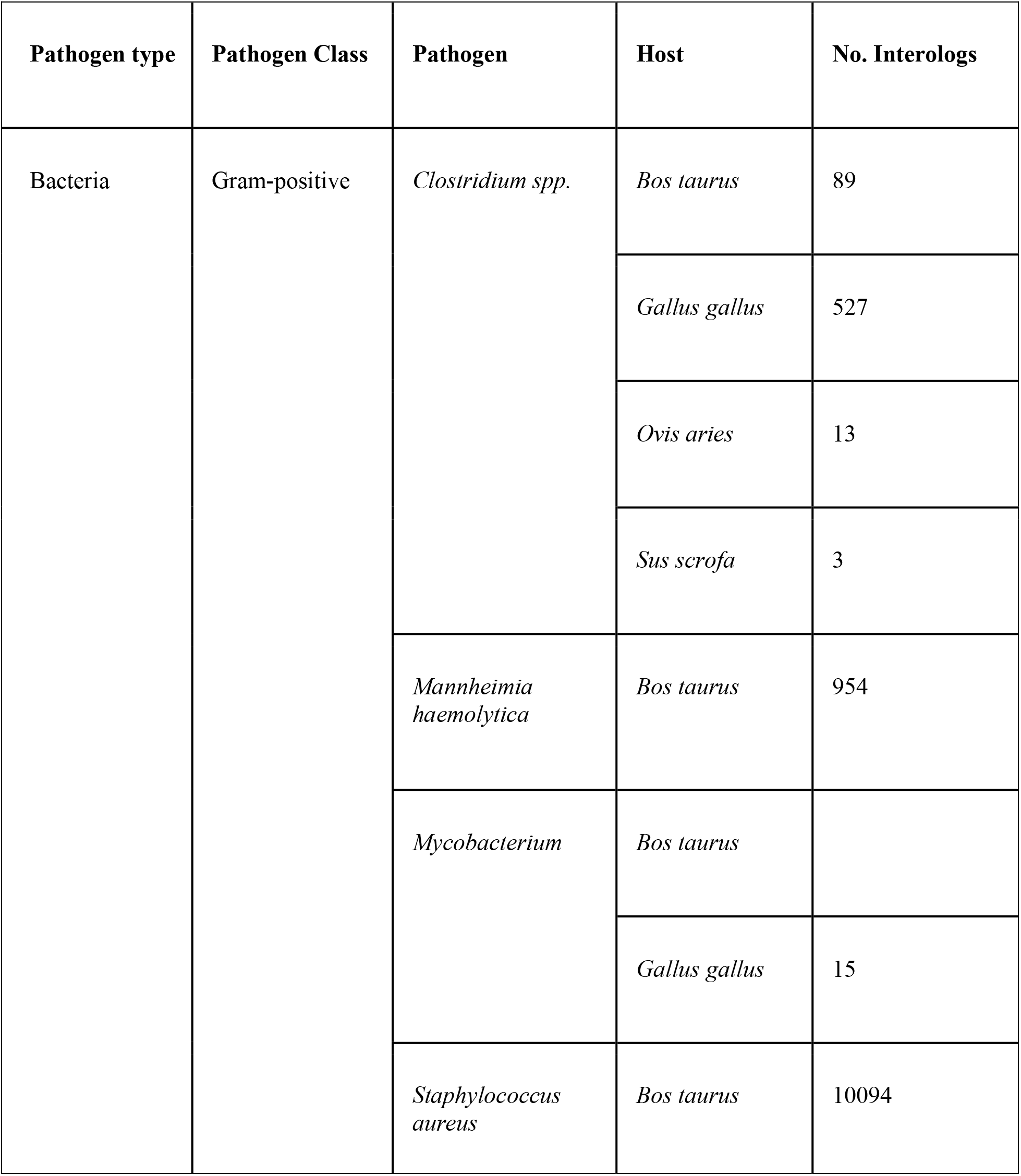
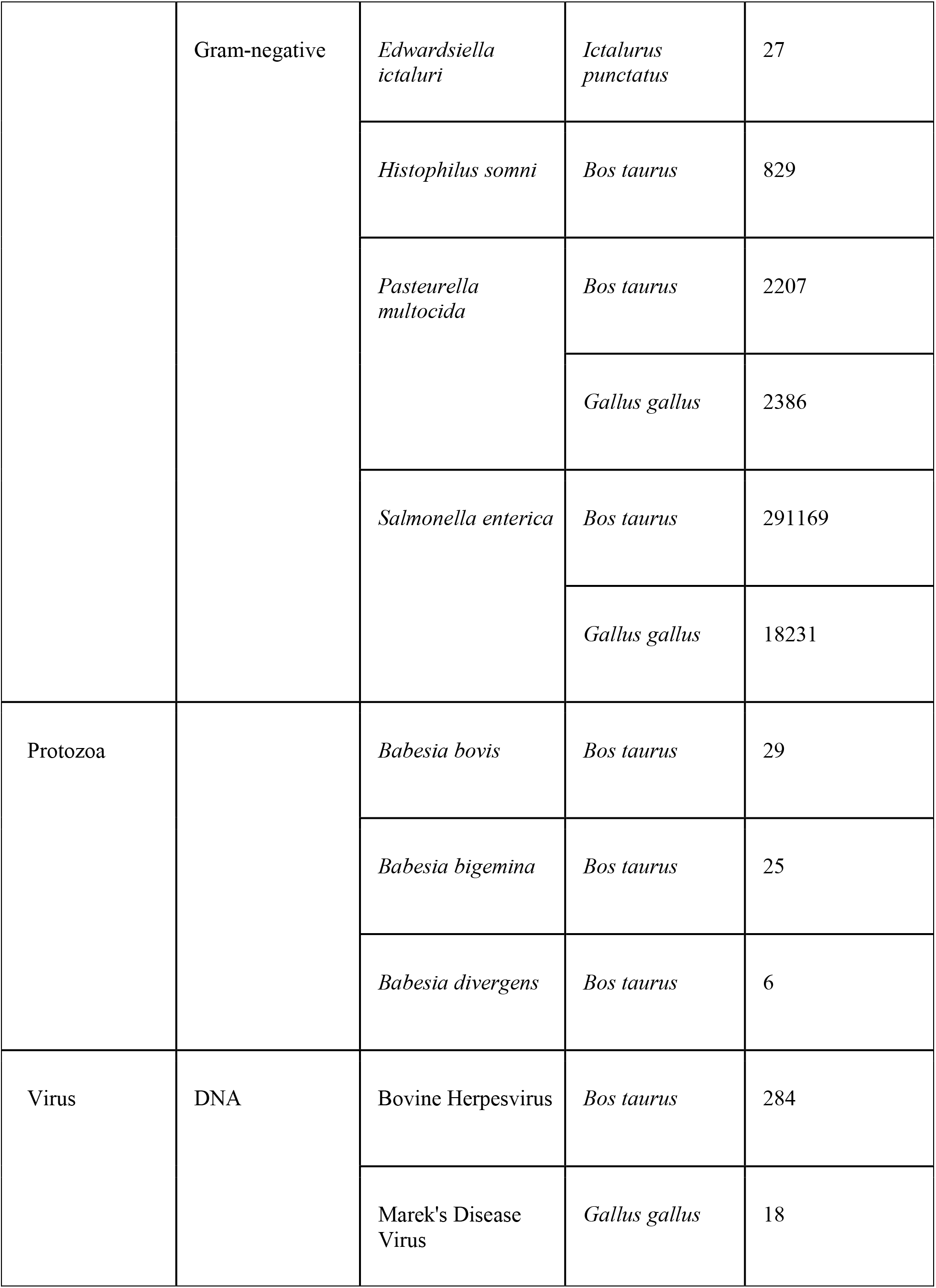
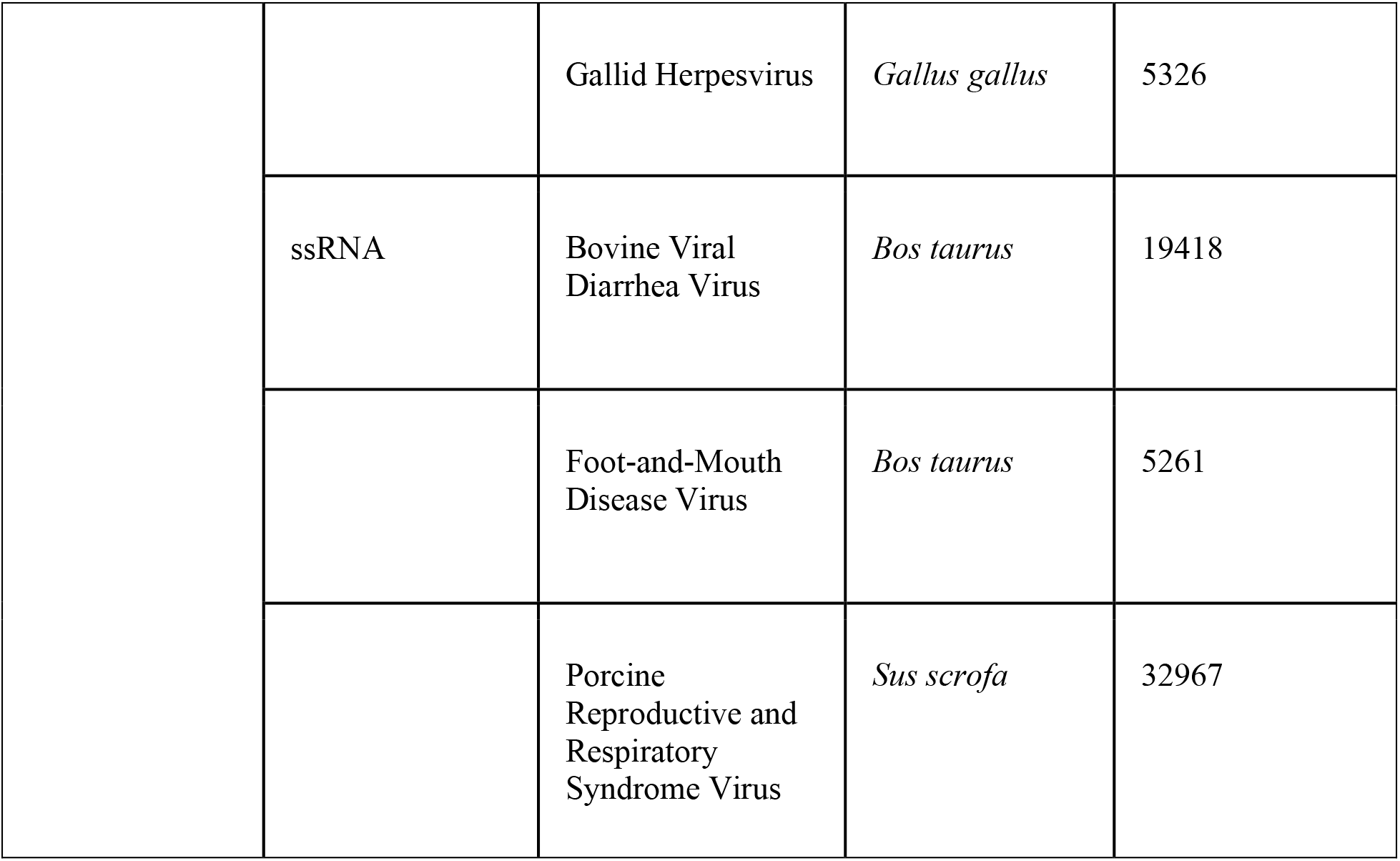
Interolog based prediction of host-pathogen interactions for agricultural host-pathogen systems.

### Interolog prediction

Predicting interolog pairs requires information regarding the orthology or homology for each protein involved in protein-protein interactions (Figure 1). For HPI pairs this requires identifying both functionally conserved host proteins and pathogen proteins. Since the hosts we analyzed are vertebrates, we initially examined the frequency of experimental vertebrate host data in the PSICQUIC Registry (23), identifying 11,371 host proteins that meet this criterion (as of July 2019). Not surprisingly most curated interactions were based upon human; for mammalian hosts we used Ensembl (Release 97) orthology to human, mouse, rat, bovine and horse, while for chicken we used orthology to human, mouse and rat as the basis of transfer between host proteins. For bovine, horse, pig and chicken host proteins we also identified orthologs via the HGNC HCOP tool (24), which includes orthology predictions from multiple sources. Only host proteins with strict 1:1 orthology and confidence = 1 were used. Since HCOP analysis does not identify orthologs to Channel catfish (*Ictalurus punctatus*, taxon ID:7998) in any species, using Channel catfish protein sequences from UniProt (version 2019-06), orthologs were determined using InParanoid ver 4.1 (25), and Blast version 2.5.0. Blast search parameters included BLOSUM62 matrix with -soft_masking, two_pass and bootstrap options, and using only one2one orthologs.

**Figure 1.** Inferring host-pathogen protein-protein interologs. A protein-protein interaction in one species can be inferred in another species if orthologs of both interacting proteins are identified in the second species. Host-pathogen interactions are a special instance of interologs which involve multiple species. We note that in many cases orthologous pathogen proteins are not identifiable, and in this case homology is used to transfer the interacting pathogen protein.

Reference pathogen protein sets were prepared by identifying pathogen proteins that had an experimentally validated HPI in PSICQUIC, and subsets of these HPI were used to generate bacterial, protozoa and viral sequence databases. Homologous pathogen proteins were identified by using proteins from our pathogens of interest (Table 1) as the queries in BlastP (ver 2.5.0) searches against the reference pathogen proteins. Search parameters used for conducting Blastp against each class of pathogen were determined after manually reviewing 100 randomly selected sequences. Based upon this manual review, we identified the following BlastP parameters: BLOSUM62 matrix, % identity > 74, % coverage > 50, Eval < 6e-180 for bacterial proteins and Eval < 2e-161 for viral and parasite proteins.

Once host orthologs and pathogen homologs are determined for a given host-pathogen system of interest, the results are compared against the original set of experimentally verified HPI to identify interolog pairs.

### Assigning confidence indicators to predicted HPI

Since our predicted HPIs are based solely on sequence similarity, we also assigned confidence indicators based upon biological features commonly found in host-pathogen protein interactions, to help evaluate the quality of the predictions. We consider predicted HPI that include these additional features to be more likely to occur in vivo. These confidence indicators (denoted by a single letter classification; Table 2) are provided to indicate a qualitative assessment of interolog predictions. The biological features of predicted HPI that are used to assign confidence are described below.

**Table 2.** Confidence indicators for interolog based HPI. At HPIDB all predicted interactions are designated with the classifier “I” (for interolog), and additional confidence classifiers are added upon these biological categories.

| Classification | Biological Relevance | Evidence |
| --- | --- | --- |
| L | host and pathogen proteins are co-localized | GO Cellular Component |
| P | host protein is involved in replication and host cell defenses | GO Biological Process |
| M | interologs contains secreted or membrane bacterial proteins | GO Cellular Component & Biological Process |
| D | pathogen protein has domains that commonly interact with host proteins | InterPro |

#### Co-localization

Proteins that are in the same cellular compartment are more likely to interact (26,27). We used Gene Ontology (GO) Cellular Component information to determine whether the predicted HPI protein pair is assigned to the same subcellular location. GO Cellular Component annotations were obtained from the GOA UniProt gene association file (2019-06 released July 3 2019). Since the GO is a hierarchical structure and annotations may occur at different “levels” in this structure, we used custom scripts to check that assigned host and pathogen GO Cellular Component terms were on the same branch of the ontology structure. The exception to this rule is the viral proteins: in the GO, viral proteins are annotated to terms preceded with “host cell” (e.g., a virus protein annotated to GO:0033644 *host cell membrane* would match a host protein annotated to GO:0016020 *membrane*). Predicted HPI pairs that meet this criterion (i.e. belonging to the same GO Cellular Component) are assigned the letter ‘L’ in the confidence assessment.

#### Secreted and membrane proteins

Bacterial proteins that interact with host are often external, secreted or membrane proteins (28,29). To determine whether the bacterial proteins from predicted HPI belonged to this category, we evaluated the GO associated with these proteins for the following GO terms (or their child terms): GO:0005576 *extracellular region*, GO:0005615 *extracellular space*, GO:0016020 *membrane*, GO:0009274 *peptidoglycan-based cell wall*, GO:0009289 *pilus*, GO:0030115 *S-layer*, GO:0043657 *host cell* and GO:0009288 *bacterial-type flagellum*. In addition, this category also includes any pathogen proteins directly annotated to GO terms that describe host-pathogen protein interactions (or their children): GO:0051701 *interaction with host*, GO:0052047 *interaction with other organism via secreted substance involved in symbiotic interaction*, and GO:0046789 *host cell surface receptor binding*. Predicted interologs that meet this criterion are assigned the letter ‘M’ in the confidence assessment.

#### Infection perturbed host processes

To establish infection, pathogens target specific host proteins involved in host innate immune defense against infections and modulate processes such as the degradation of proteins by the proteasome, ubiquitination of proteins, apoptosis and cell cycle etc. (15,30–32). HPIs containing host proteins involved in replication and host cell defenses, based on GO Biological process are assigned the letter ‘P’. These include interologs where the host protein is annotated to the following GO terms (or their children): GO:0006915 *apoptosis*, GO:0007049 *cell cycle*, GO:0043248 *proteasome assembly*, GO:0031144 *proteasome localization*, GO:0016567 *protein ubiquitination*, GO:0045335 *phagocytic vesicle* and GO:0045087 *innate immune response*. Predicted interologs that meet this criterion are assigned the letter ‘P’ in the confidence assessment.

#### Interacting domains

HPIs are frequently mediated by conserved protein motifs and domains, and computational approaches utilize this knowledge to identify host-pathogen interactions (33). We also utilize this approach, using a combination of InterproScan (34) to identify motifs of proteins known to be involved in host-pathogen interactions and 3did (35) to predict interacting domains between host and pathogen proteins. All host interolog and pathogen interolog sequences were analyzed using InterproScan v5.35-74.0 and 3did (ver. 2019_01) with the Pfam-hmm file (Pfam version 32.0). The 3did analysis utilized hmmer-3.1b2 to find Pfam domains for host interolog and pathogen interolog sequences, parsing hmmsearch results and 3DID interacting domains to find Pfam 3DID evidence. The details of the motifs we used, their function and the protein datasets we searched to identify these motifs (e.g., host or pathogen) are specified in Table 3. Predicted HPI that meet this criterion are assigned the letter ‘D’ in the confidence assessment.

**Table 3.** Domains used to apply confidence indicator D to predicted host-pathogen interactions.

| **Motif/Domain** | **InterPro ID** | **Function** | **References** | **Protein set identified with motif** |
| --- | --- | --- | --- | --- |
| BAR | IPR004148<br>BAR domain | endocytosis and intracellular transport | 26118995,<br>27265768 | host proteins in host-pathogen interactions |
| CRIB | IPR000095<br>CRIB domain | conserved region of a GTPase binding domain found in downstream effectors of the small GTPases CDC42 and RAC | 19709124<br>12046591<br>22145094 | host proteins in host-bacterial interactions |
| PDZ | IPR001478<br>PDZ domain | localize cellular elements and regulate cellular pathways | 21775458,<br>21517912 | host proteins in host-pathogen interactions |
| Rho GTPase-activating protein domain | IPR000198<br>Rho GTPase-activating protein domain | transduce signals from plasma-membrane receptors | 25203748,<br>11680784,<br>24691164 | host proteins in host-pathogen interactions |
| SH2 | IPR000980<br>SH2 domain | mediate protein-protein interactions; control cytoplasmic signaling | 7859737,<br>24175231,<br>19380118 | host proteins in host-pathogen interactions |
| SH3 | IPR001452<br>SH3 domain | mediate protein-protein interactions; control cytoplasmic signaling | 7859737, 24175231, 19366662, 24600591 | host proteins in host-pathogen interactions |
| SNARE | IPR010989<br>SNARE domain | membrane fusion | 21178463, 21680744 | pathogen proteins |

### Assessment of predicted interolog HPI data

To assess the impact of our predicted host-pathogen interactions we analyzed (a) the proportion of host proteins that are hubs or bottlenecks and (b) the impact of adding these interologs to a network analysis. Hubs and bottlenecks were identified by determining network centralities for degree (connectivity) and betweenness centrality (bottlenecks). We used STRING 11.0 (22) to obtain 11, 259, 366 bovine interactions. These were filtered to obtain high quality interactions by setting a confidence score of ≥ 700. Mapping the high confidence Ensembl protein interaction accessions to UniProtKB protein accessions generated a set with 19, 304 nodes with 526, 261 edges/interactions. This data set was analyzed using Cytoscape 3.7.2 (36) NetworkAnalyzer (37) to calculate network centrality scores for degree and betweenness, and the top 20% of both degree and betweenness centrality scored bovine proteins were selected as hubs and bottlenecks, respectively. This set of bovine hub and bottleneck proteins was used to identify predicted HPIs where the pathogen protein targeted bovine hubs and bottlenecks.

We also assessed the contribution of interologs to network analysis using bovine-BVDV as an example. Analysis of the PSICQUIC Registry version 1.4.11 (23) identified existing bovine-BVDV host-pathogen interactions, and this set provided the initial HPI dataset of network analysis. We supplemented these HPI with (a) all bovine-BVDV interologs and (b) bovine-BVDV interologs with confidence indicators to create comprehensive and high confidence HPI data sets, respectively. Each data set was imported into CytoScape 3.7.2 and analyzed to create an undirected network. NetworkAnalyzer was used to analyze network statistics and networks were visualized using this tool to map node size to betweenness centrality, assigning low values to small node sizes and dark node color to low values. GO enrichment was done using the Panther Overrepresentation tool (38) (Released 20200407) with all *Bos taurus* genes as the reference set, GO annotations as at 2020-03-23, and a Fisher’s test with Bonferroni correction.

## RESULTS AND DISCUSSION

### Predicted interactions

The agriculturally related pathogen-host pairs we used for HPI prediction were selected based upon funded animal health projects and available HPI data, and these pairs are summarized in Table 1. A total of 389,878 predicted HPIs are identified for these 23 host-pathogen pairs. Only 76 of these predicted interactions are previously annotated by interaction databases, and this reflects the dearth of large-scale curation of agricultural HPIs. Interactions were predicted from a combined set of 6,364 pathogen proteins from 40 pathogen strains (with information on species, sub-species and strains included). Likewise, 11,371 host proteins from five host species were investigated to identify these interologs. Not surprisingly, host-pathogen pairs with sequence similarity to well curated human-pathogen pairs had the most predicted data. For example, bacterial pathogens made up 84% of the pathogen reference set and produced 82% of the total predicted HPI. Likewise, bovine proteins match well with human proteins, and host-pathogen pairs with a bovine host make up 85% of these predicted HPI.

The interolog based predictions are a first-pass set of HPIs for agricultural pathogens. However, there are some limitations to this method. First, the paucity of experimentally verified interaction data which is the basis for these predictions hinders this approach. The majority of experimental HPIs available in bioinformatics databases are for human pathogens, which limits interolog predictions to related host-pathogen pairs. In addition, lack of protein information and curation for some types of pathogens (e.g., protozoa species) resulted in very few pathogen protein matches for interolog prediction, particularly using high stringency parameters required for Blast. Our ability to identify host-pathogen interologs will only improve as the annotation of pathogen proteins improves, and as more diverse HPI data is curated. We note that the prediction of interologs has helped us to identify potential host-pathogen pairs to target for curation, with the goal of also improving interolog prediction.

Traditionally, assessment of prediction requires calculation of measures such as accuracy and precision. However, these measures require information regarding both true positives and true negatives. While experimentally validated interactions can serve as true positives, this set is necessarily incomplete; worse still, negative interactions are not reported. Therefore, we evaluated interolog predictions by assessing their likelihood and biological relevance. To interact, proteins should co-localize and are expected to be involved in the same cellular processes and pathways. For example, pathogens target processes to alter host immune responses (39). Additionally, it is well documented that pathogen proteins contain specific interacting motifs critical for host interactions (40–43). We used GO Cellular Component and Biological Process annotations (44) and functional motifs data from InteroPro (45) to identify predicted HPI that met these criteria. In each case we assigned predicted HPI with a single letter code based on the rule set we developed by biological assessment (Table 2). When we applied these assessments to our predicted HPI set, 39.5% met at least one of these criteria (Figure 2A), with interactions containing host proteins involved in replication and host cell defenses (“P”) the most assigned rule (Figure 2B). We note that assessment rules are based on the current understanding of how host and pathogen proteins interact (e.g., ability to co-localize, common interacting domains and commonly targeted processes). As our knowledge of host-protein interactions increases, it will be possible to add and edit additional rules to improve the assessment of predicted HPI. We also predict that the use of the interolog data for modeling and experimental validation will provide feedback that will further improve biological assessment.

**Figure 2.** Applying confidence classifiers to predicted interactions. (A) The number of confidence rules assigned to predicted interactions. (B) The number of confidence rules assigned to predicted interactions by category.

### Assessment of predicted HPI

Ultimately, the value of HPIs lies in their use for modeling interactions, most commonly using network analyses. These network analyses include assessment of protein connectivity (i.e., the number of interactions), and perturbed cellular pathways and functions (46,47). While our interolog based approach identifies a number of HPI for host-pathogen systems, we also assess whether predicted interactions inform our knowledge of host-pathogen interactions during infection. First, we assess the connectivity of networks that include predicted HPIs to examine highly connected proteins. Then we generate host-pathogen networks both with and without our predicted interologs and evaluate the information contained in these networks.

#### Connectivity of predicted HPIs

Pathogen proteins frequently target host proteins that serve as hubs and bottlenecks (39) to affect essential cellular processes. We used network analysis of bovine proteins to identify bovine hubs (proteins with high connectivity) and bottlenecks (centralness) (46,47) and determine the frequency of these proteins in our predicted host-pathogen interactions. We identified 603 bovine hub and bottleneck proteins. These same proteins were involved in 29% or our predicted bovine interologs, and this proportion increases to 38% among predicted bovine interologs that also contain at least one confidence indicator (Figure 3). We note that the proportion of hub or bottleneck proteins targeted by pathogens also varies based on the pathogen type, with viruses - representing intracellular pathogens - not unexpectedly interacting with the highest proportion of host hubs and bottlenecks.

**Figure 3.** Proportion of predicted pathogen interactions with bovine hub and bottleneck proteins. Subsets of bovine interologs that include at least one confidence indicator (L, P, M or D) are also indicated.

Analysis of human hubs and bottlenecks identified that 49% of proteins serve as both a hub and a bottleneck (48). Our analysis of bovine proteins supported this, with 50% of bovine hubs also identified as bottlenecks. Bovine proteins that act as hubs that are highly represented in our bacterial interologs include NFKB1, von Willebrand factor and catenin beta-1. NFKB1 controls programmed cell death and is known to be targeted by multiple pathogenic bacteria (49) (Table 4). von Willebrand factor and catenin beta-1 both play a role in inflammation and are targeted by multiple bacterial species (50–52). STAT6, another protein that is highly represented in our bovine interolog set, was identified as a bottleneck. STAT6 also plays a role in inflammation (53). GO enrichment analysis of bovine hubs and bottlenecks identified from our bacterial interolog data set demonstrates that these host proteins are involved in *negative regulation of epithelial cell differentiation, translational elongation* and *cellular response* processes (Table 5). These functions represent interactions of pathogenic bacteria with epithelial surfaces, translation and responses to bacteria and secreted bacterial peptides.

**Table 4.**
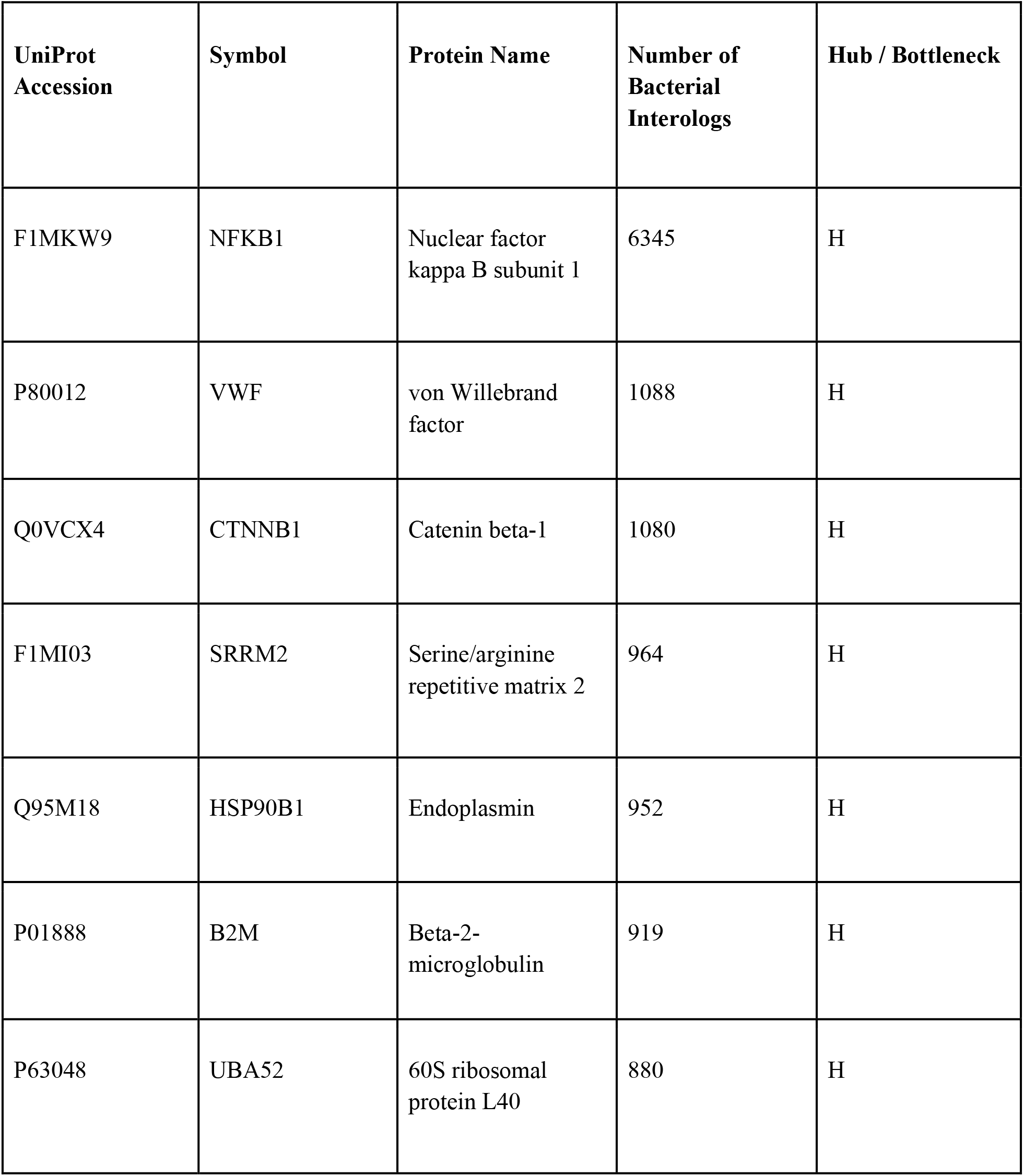
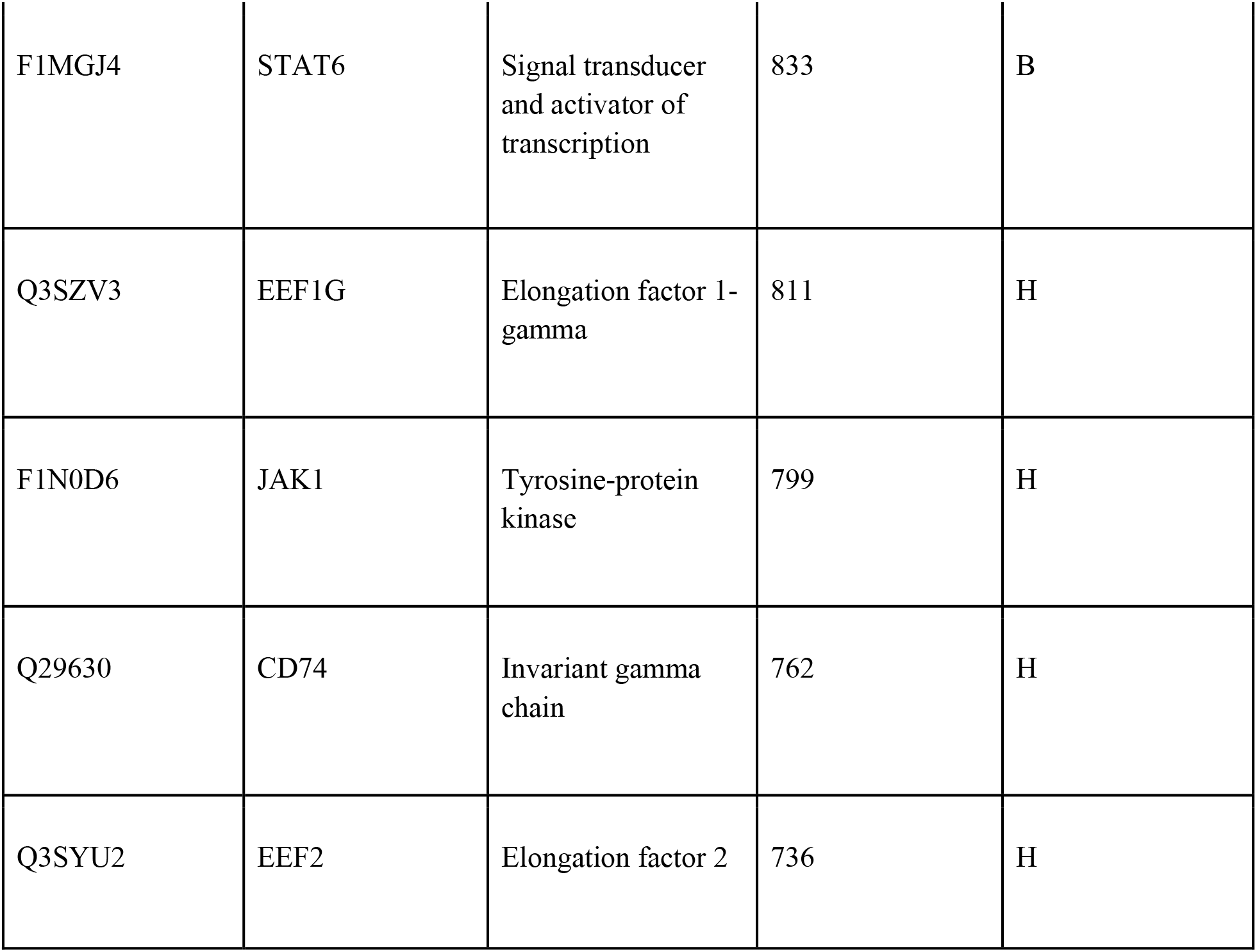
Bovine hub and bottleneck proteins which are highly represented in predicted interolog based HPI. Hubs and bottlenecks are indicated with a H or B, respectively.

**Table 5.** Enriched functions of bovine hub and bottleneck proteins which are highly represented in predicted interolog based HPI. Based upon the PANTHER Overrepresentation Test (Released 20200407) using Fisher’s Exact test and Bonferroni multiple testing correction. GO Biological Process annotations from bovine proteins were used as the reference set.

| GO biological process | Fold Enrichment | +/- | p-value |
| --- | --- | --- | --- |
| negative regulation of epithelial cell differentiation | > 100 | + | 1.37E-04 |
| translational elongation | 99.58 | + | 3.37E-02 |
| response to peptide | 29.87 | + | 2.77E+00 |
| cellular response to organonitrogen compound | 17.84 | + | 5.46E-03 |
| response to hormone | 14.81 | + | 1.58E-02 |
| response to drug | 13.89 | + | 2.77E-03 |
| cellular response to oxygen-containing compound | 11.57 | + | 9.37E-03 |

As hypothesized, the predicted HPIs are enriched for host proteins that serve as hubs and bottlenecks. Moreover, these host proteins themselves have functional roles that are likely to be targeted by pathogens. Taken together, these characteristics support the usefulness of the interolog approach in predicting HPIs to support network analysis of host-pathogen pairs where there is little manual curation available to support network analysis. We next assess the use of these interologs to determine if they could provide information about possible intervention targets for disease control.

#### Adding interolog data to network analysis: bovine-BVDV network analysis

To determine if interologs provide insights into network analysis of host-pathogen pairs, we compared bovine-BVDV interactions from existing, manually curated HPI (referred to as “existing”), this data supplemented with all predicted interologs (“interologs”) and the PSICQUIC interactions supplemented only with interologs annotated with confidence indicators (“high confidence”). The results of these network analysis are shown in Figure 4. The bovine-BVDV network based upon existing, manually curated interactions (Figure 4A) reflects well studied BVDV E2 and NS3 proteins, showing their interactions with host proteins but little else. When interologs are added to the network (Figure 4B & 4C), other viral proteins are represented in the network and the number of connected nodes increases. Notably, the addition of interologs to the network analysis identified CD46 and TXN2 host proteins which are central to the network. CD46 (identified in both networks that include interologs) inactivates complement and induces Treg1 cells which suppress immune responses (54,55); this protein is also the BVDV receptor (56). TXN2 (thioredoxin 2) interacts with multiple BVDV proteins in the “all interolog” network and has high centrality, while in the high confidence network is less central and directly linked only to BVDV E2. TXN2 protects against oxidant-induced apoptosis and has been demonstrated to inhibit viral replication of another Pestivirus, Classical Swine Fever Virus, by binding to its E2 protein (57). This demonstrates that the utilization of predicted HPIs supports informative host-pathogen network analysis.

**Figure 4.** Bovine-BVDV networks. (A) Based upon existing HPIs from PSICQUIC. (B) Existing interactions combined with all predicted bovine-BVDV interologs. (C) Existing interactions combined with all predicted bovine-BVDV interologs that are assigned a confidence indicator. Proteins are represented by circles and interactions by connecting lines. Larger, red circles indicate hubs and bottlenecks, which in most cases are viral proteins. BVDV proteins are also outlined in red.

In addition to the networks themselves, we also looked at GO enrichment of host proteins represented in these networks. While there were no statistically significant process terms enriched from the existing set of host proteins, enriched biological processes for all interologs and high confidence interactions are in concordance (Table 6). Processes such as *regulation of protein stability, positive regulation of cellular protein localization* and *negative regulation of cell death* reflect RNA virus replication strategies. We note that while both data sets containing interologs had similar networks, several functional differences occur. Metabolic process terms and *symbiotic process* are enriched exclusively in the high confidence data set, while these terms are not identified in the “all interolog” data set. Viruses rely on host cell machinery for synthesis of macromolecules required for viral replication (58) and the *symbiotic process* term is the parent GO term that encapsulates specific viral processes. This may indicate that specifying the set of predicted HPIs with confidence indicators will provide more relevant information, although this conclusion is tempered by the knowledge that the number of high confidence bovine-BVDV interologs was high enough to be informative, which may not be the case in other host-pathogen pairs.

**Table 6.** Enriched functions from bovine-BVDV host proteins. Based upon the PANTHER Overrepresentation Test (Released 20200407) using Fisher’s Exact test and Bonferroni multiple testing correction. GO Biological Process annotations from bovine proteins were used as the reference set.

| GO biological process (GO:ID) | all interologs |  | high confidence |  |
| --- | --- | --- | --- | --- |
|  | fold enrichment | p-value | fold enrichment | p-value |
| symbiotic process (GO:0044403) | - | - | 11.92 | 1.87E-02 |
| regulation of protein stability (GO:0031647) | 8.94 | 5.27E-04 | 11.56 | 4.27E-03 |
| positive regulation of cellular protein localization (GO:1903829) | 8.3 | 4.07E-03 | 10.34 | 4.65E-02 |
| negative regulation of cell death (GO:0060548) | 4.11 | 1.61E-02 | 6.4 | 2.55E-04 |
| regulation of phosphorylation (GO:0042325) | 3.18 | 1.18E-02 | - | - |
| organic substance catabolic process (GO:1901575) | 3.11 | 9.59E-03 | - | - |
| response to organic substance<br>(GO:0010033) | 2.76 | 3.34E-03 | 3.58 | 2.53E-02 |
| regulation of protein metabolic<br>process (GO:0051246) | - | - | 2.97 | 2.63E-02 |
| regulation of molecular function<br>(GO:0065009) | - | - | 2.87 | 4.33E-02 |
| cellular metabolic process<br>(GO:0044237) | - | - | 1.92 | 4.21E-02 |

#### Availability of data and materials

Predicted interologs are available from the HPIDB database (hpidb.igbb.msstate.edu/hpi30_interologs.html) as a downloadable MITAB file (59), can be selected for inclusion in HPIDB searches, and used for network analysis with software such as Cytoscape (37). Confidence indicators are given in the confidence column of the MITAB file (column 15). In addition, a MITAB_plus file that contains 11 additional columns with information about the interacting proteins’ taxon category and sequence etc. is provided. All files include the confidence assessors discussed earlier and all interolog predictions are marked with an “I” to distinguish these HPI from experimentally derived interactions in the HPIDB.

## CONCLUSIONS

We extend the interolog protein-protein prediction method by applying it to predict host-pathogen interactions and utilizing a biology-based rule system to assign confidence to these predictions. As expected, extending prediction of HPI to additional host-pathogen pairs is limited by the availability of high confidence, manually curated HPI data. We note that this example of extending high-quality, manual biocuration to related species increases the value of these critical manual biocuration efforts (60). Confidence indicators are included in the predicted HPI datasets and are given letter codes so that they cannot be mistaken for a quantitative assessment. Moreover, we demonstrate that host proteins in the predicted HPI are enriched for hubs and bottlenecks, which correlates with our understanding of proteins targeted by pathogens. Likewise, when we included our predicted interologs in a network analysis, the resulting analyses recapitulates biological information about pathogenesis in a way that is not possible when only previously existing HPI interactions are used, demonstrating that predicted HPI can contribute to a better understanding of infections of relevance to US agriculture.

## DECLARATIONS

### Ethics approval and consent to participate

Not applicable.

### Consent for publication

Not applicable.

### Authors’ contributions

All authors contributed to the conception, design and analysis of this work. MA interpreted data, did network analyses and completed the manuscript draft. CG developed scripts to collect data and completed computational analyses. BN and FM developed confidence rules, interpreted data and contributed to the preparation of this manuscript. All authors have read and approved this manuscript.

## Acknowledgements

Not applicable.

